# Engineered acetoacetate-inducible whole-cell biosensors based on the AtoSC two-component system

**DOI:** 10.1101/035972

**Authors:** Jack W. Rutter, Linda Dekker, Alex J. H. Fedorec, David T. Gonzales, Ke Yan Wen, Lewis E. S. Tanner, Emma Donovan, Tanel Ozdemir, Geraint Thomas, Chris P. Barnes

## Abstract

Whole-cell biosensors hold potential in a variety of industrial, medical and environmental applications. These biosensors can be constructed through the repurposing of bacterial sensing mechanisms, including the common two-component system. Here we report on the construction of a range of novel biosensors that are sensitive to acetoacetate, a molecule that plays a number of roles in human health and biology. These biosensors are based on the AtoSC two-component system. An ODE model to describe the action of the AtoSC two-component system was developed and sensitivity analysis of this model used to help inform biosensor design. The final collection of biosensors constructed displayed a range of switching behaviours, at physiologically relevant acetoacetate concentrations and can operate in several *Escherichia coli* host strains. It is envisaged that these biosensor strains will offer an alternative to currently available commercial strip tests and, in future, may be adopted for more complex *in vivo* or industrial monitoring applications.

## Introduction

Bacterial cells have evolved the ability to detect and respond to environmental cues. The repurposing of these natural abilities to create engineered biosensor strains has become a major area of research within synthetic biology. Whole-cell biosensors, which contain this sensing machinery within a single living cell, offer a number of advantages over more conventional sensing technologies including portability, self-replication, sensing of the bioavailable fraction (rather than total concentration) and low-cost of production[1, 2, 3, 4]. To date, whole-cell biosensors have been developed for the detection of metals in environmental samples (mercury[5], arsenic[6], lead[7]), for the monitoring of metabolites during bioproduction (lactate[8]) and in medical applications for the monitoring of clinically relevant biomarkers [9, 10, 11, 12, 13], exploiting the ability of bacteria to reach inaccessible areas of the host *in vivo*[4]. Whole-cell biosensors can be based on a number of bacterial sensing mechanisms. These include transcription-factor mechanisms, one-component systems, two-component systems (TCSs), extracytoplasmic function sigma factors, and other environment responsive mechanisms[1, 14].

TCSs are one of the most common bacterial sensing mechanisms, with many examples found within bacterial genomes[15, 16]. *Escherichia coli* alone are reported to have 32 TCSs[15]. TCSs typically consist of a membrane-bound histidine kinase (HK) that detects a specific signal. This signal activates kinase activity and causes autophosphorylation of a conserved histidine residue. The phosphoryl group is then transferred to an aspartate residue on the response regulator (RR)[16]. Usually RRs are transcriptional regulators capable of activating or repressing expression from a specific promoter[17], thereby controlling gene expression in response to an external input. TCSs respond to a wide range of input signals, including quorum-sensing molecules[18], certain wavelengths of light[19], physical contact[20] and oxidative stress[21]. Previously reported TCS whole-cell biosensors have been developed for tetrathionate[9], thiosulfate[10], nitrate[11] and aspartate[12].

The AtoSC TCS is sensitive to acetoacetate and within *E. coli* plays a role in short-chain fatty acid (SCFA) metabolism, motility and poly-(R)-3-hydroxybutyrate synthesis [22, 23, 24, 25]. As shown in Figure 1, acetoacetate is detected by the membrane-bound AtoS histidine kinase (AtoS HK). This then goes on to phosphorylate the AtoC response regulator (AtoC RR). The phosphorylated AtoC RR then induces transcription from the Pato promoter, which subsequently expresses the *atoDAEB* operon.

**Figure 1:**
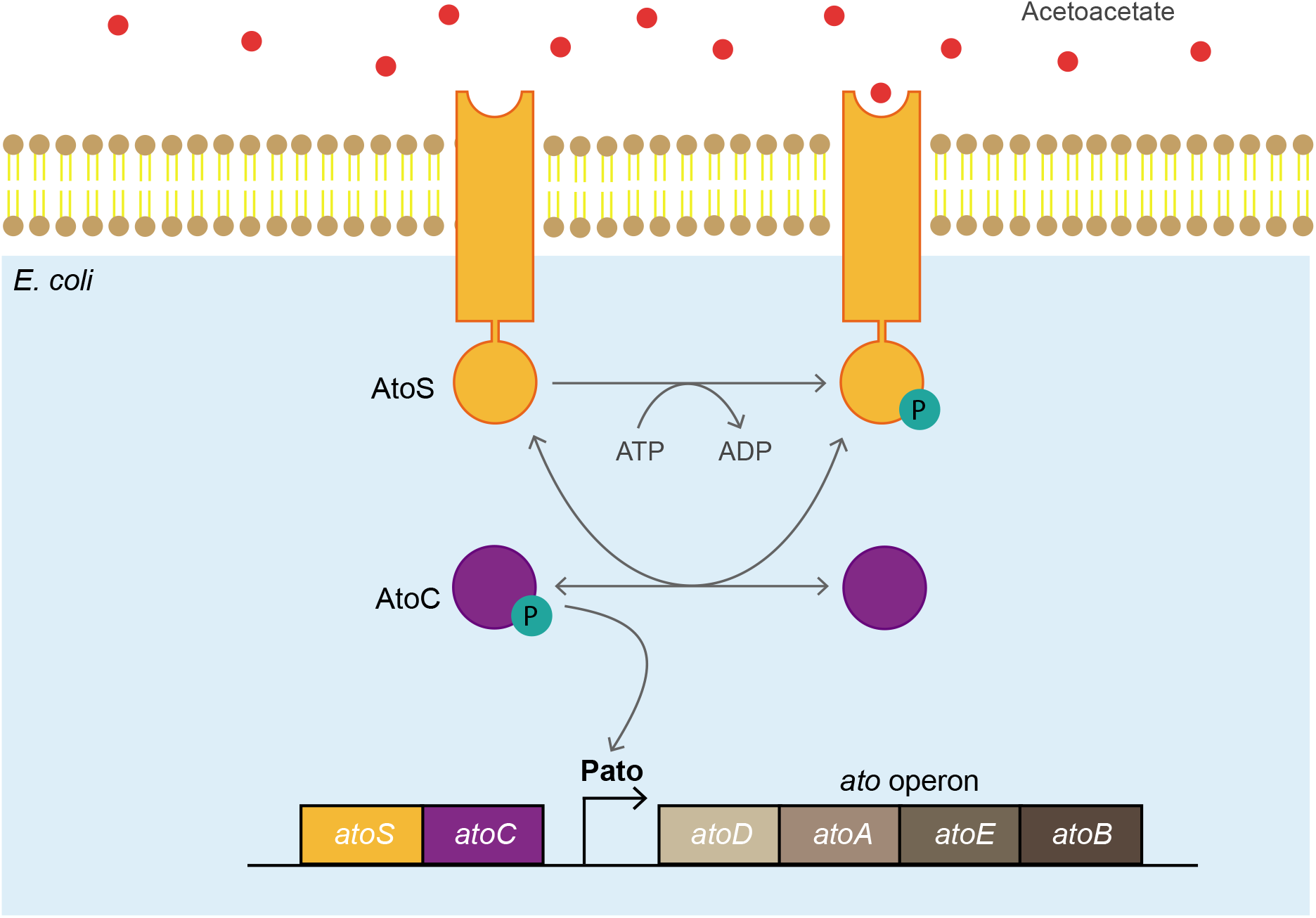
Layout of the AtoSC two-component system and *atoDAEB* operon. The AtoS histidine kinase, within the *E. coli* inner membrane, autophosphorylates in the presence of acetoacetate. The phosphate group is then transferred to the AtoC response regulator, which triggers expression from the Pato promoter.

Acetoacetate is an extremely important metabolite in mammalian metabolism and energy regulation. It is one of three ketone bodies, alongside *#x03B2;*-hydroxybutyrate and the less abundant ace-tone, which can serve as an alternative energy source in the body during periods of low glucose availability[26]. Typically ketone bodies are found at sub-millimolar concentrations within the blood; however, these levels can become elevated during periods of extended starvation or intense exercise[27]. Ketone bodies have been linked to protective effects on the neural system, which has led to the use of ketogenic diets as a method for preventing epileptic seizures[28]. Another study highlighted the role ketone bodies play in stem cell differentiation and homeostasis within the intestines, through the action of *#x03B2;*-hydroxybutyrate[29]. Human mesenchymal stem cells have been shown to have a preference for acetoacetate as an energy-yielding substrate; leading to suggestions that acetoacetate could be added to mesenchymal stem cell culture medium[30]. Acetoacetate has also been shown to act as a signalling molecule for muscle regeneration and can help restore muscle function in a mouse model of muscular wasting[31]. However, sustained periods of increased ketone body concentrations within humans can be a sign of several pathological states. These include salicylate poisoning, alcoholic ketoacidosis and diabetic ketoacidosis[27]. During extreme cases of diabetic ketoacidosis ketone body concentrations can reach 20 mM and above. If left untreated, this may lead to complications such as a cerebral edema[32, 26]. As such, ketone body levels (particularly acetoacetate and *β*-hydroxybutyrate) are monitored regularly in patients with diabetes. Alongside their role in diabetes, ketone bodies are also measured in the blood of cows. During pregnancy, cows become susceptible to ketosis, a condition that adversely affects the health and milk production of dairy cattle[33].

It is clear therefore that methods of measuring ketone body levels may find use in a range of biomedical or agricultural applications. Currently, strip tests that use a sodium-nitroprusside reaction, or ketone meters, can be used to monitor ketone levels in urine and blood[34]. Although these offer cheap monitoring systems for patients, they are only semi quantitative, and there are a number of biomedical, scientific and industrial applications where they are unsuitable. For example, monitoring the production of acetoacetate by engineered *E. coli*[35], exploring how the human microbiota utilises ketone bodies [36] and investigating host-microbiota interactions in model systems *in vivo* [37], could all use whole-cell biosensors to report on acetotacetate concentration.

Here, we report the construction of *E. coli* whole-cell biosensors that can be used to detect and report on the presence of acetoacetate. The whole-cell biosensors presented here are based on the AtoSC TCS, found within *E. coli*. We develop an ODE model that attempts to capture the action of the AtoSC TCS and use sensitivity analysis of this model to guide the design of an array of acetoacetate-inducible biosensors. The final biosensors display a diverse range of output responses and may be employed in the future for a variety of sensing applications.

## Materials and Methods

### AtoSC plasmid construction

The plasmids within this study were constructed using the CIDAR MoClo assembly method[38]. The oligonucleotides and plasmids designed in this study are given in SI Tables S1 and S2, respectively, alongside details of their construction.

### AtoSC biosensor host strains

A range of host strains were used as chassis for the AtoSC biosensor plasmids. The BW25113 (*atoSC*^+^), JW2213 (*atoS*^−^) and JW2214 (*atoC*^−^) strains were purchased from the Keio collection (Dharmacon Horizon, Cambridge, UK). The BW28878 (*atoSC*^−^) double-knockout strain was provided by Professor Kyriakidis (University of Thessaloniki) and *E. coli* Nissle 1917 by Professor Henderson (University of Birmingham). Competent NEB5*α* cells were purchased commercially (New England Biolabs, MA, USA). All strains are given in SI Table S2.

### AtoSC ODE modelling and sensitivity analysis

An ODE model was constructed to describe the AtoSC whole-cell biosensors behaviour (Figure 2A). This modelling was performed in Python (all code to reproduce these results can be found on zenodo). The equations and parameter bounds used for this model are given in SI section 2. Sensitivity analysis was performed based on the SA.lib python package, using the Morris method[39].

**Figure 2:**
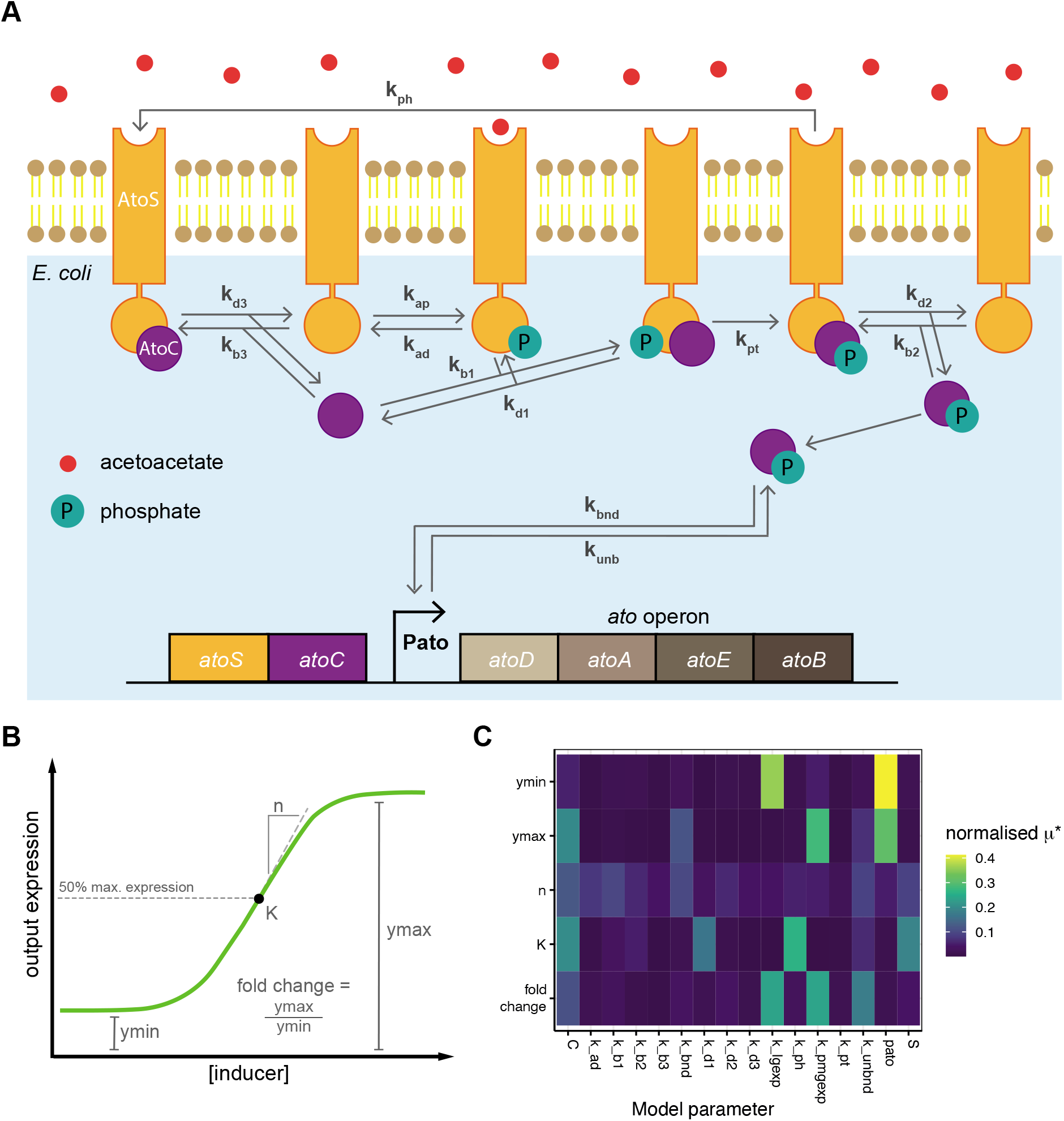
Results of the sensitivity analysis conducted on the AtoSC model. (**A**) Layout of the AtoSC model, illustrating the parameters varied during sensitivity analysis of the system. (**B**) Illustration of the different aspects of biosensor performance for which sensitivity analysis was performed, ymin and ymax represent *f^min^* and *f^max^*, respectively. *K* represents the *K*_1/2_ of the biosensor. (**C**) Heatmap denoting the normalised μ* results of the sensitivity analysis for the AtoSC model parameters, a higher *μ\** value indicates higher parameter influence on biosensor behaviour. ‘C’ refers to AtoC RR concentration, ‘S’ to AtoS HK concentration and ‘pato’ to Pato promoter presence.

#### Growth curve assays

Overnight bacterial cultures were diluted to an approximate OD_700_ of 0.05, within fresh media. 120 *μ*L of each culture was then added to the well of a 96-well clear bottom microplate (Greiner Bio-One) with a magnetic removable lid. The cultures were incubated in a Tecan Spark plate reader for 2 hours at 37°C with shaking at 150 rpm (2 mm amplitude, double orbital) with an OD_700_ measurement taken every 30 minutes. Cultures were then induced with the relevant inducer concentration and sealed with a breathe-easy permeable membrane (Diversified Biotech). The plates were then returned to the plate reader and left to grow for a further 16 hours, with OD_700_ measurements taken every 20 minutes (the same temperature and shaking conditions were maintained). Inducers were added to the 96-well plates with the aid of an I.Dot liquid handler (Dispendix, Stuttgart, Germany). Inducers were diluted to the desired stock concentrations and then custom protocols set up within the I.Dot, using the I.Dot assay studio software, to automate addition of the correct inducer volumes. Taking the total volume in each well to 125 *μ*L.

### Dose-response Assays

Bacterial cultures, grown overnight in LB media, were diluted to an OD_700_ of approximately 0.05 in fresh media, and 190 *μ*L added to each well of a polypropylene 96-well deep-well plate (Brand, Sigma Aldrich). The plate was then sealed with an autoclaved system Duetz lid and incubated for 2 hours at 37°C, with 350 rpm shaking[40]. Inducers were then added to the desired concentration, bringing the total volume in each well to 200 *μ*L, the plate was once again sealed with an autoclaved system Duetz lid and the induced cultures incubated at 37^°^C, with 350 rpm shaking for 16 hours. Samples were taken as required for flow cytometry analysis.

### Flow cytometry analysis

Flow cytometry was performed on an Attune NxT Acoustic Focusing Cytometer, with an Attune NxT Autosampler (Thermo Fisher Scientific, UK), closely following a previously reported method[37]. In brief, one *μ*L of the appropriate sample was diluted 1:200 in sterile phosphate-buffered saline (PBS), in a shallow polystyrene U-bottom 96-well plate. An Attune NxT Autosampler was used to record 10,000 events (for each sample) with 3 wash and mixing cycles between samples. GFP was excited using a blue laser (488nm) and detected using a 530/30nm bandpass filter. Additionally, a sample of 1:300 dilution of rainbow calibration particles in sterile PBS (Spherotech, UK) was recorded allowing for the conversion of arbitrary units to molecules of equivalent fluorophore (MEF), using R commands from the FlopR package[41].

All collected flow cytometry standard (FCS) data were processed using commands from the FlopR package[41] and plotted using custom R scripts. Visualisation and curve fitting were performed in R, using the ‘ggplot2’ package and ‘nls’ fitting function[42]. GFP induction data was fit using a Hill function:

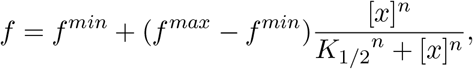

where *f* is the observed fluorescence, *f^min^* is the minimum (basal) fitted value, *f^max^* is the maximum fitted value, [*x*] is the inducer concentration, *K*_1/2_ is the threshold sensitivity and *n* is the cooperativity. Dynamic range was calculated using the following expression, based on the fitted values of *f^min^* and *f^max^*.

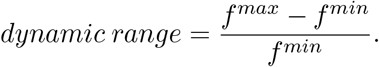

Fold change was calculated as *f^max^/f^min^.*

## Results

### Sensitivity analysis of the AtoSC TCS

An ODE model describing the action of the AtoSC TCS was developed (Figure 2A) to aid in experimental design. Morris sensitivity analysis was carried out to identify the model parameters that were predicted to have a large effect on AtoSC biosensor response. This analysis was performed for each of the major components of biosensor behaviour, *K*_1/2_, *f^min^*, *f^max^*, etc. (Figure 2B, rankings are given in SI Figure S1). The sensitivity analysis predicted several parameters would have an influence on final biosensor behaviour. The concentrations of the AtoS HK, AtoC RR proteins and the presence of the Pato promoter were all predicted to have an influence on a range of different aspects of biosensor performance (Figure 2C). In theory, these parameters are readily tuneable experimentally, through varying of plasmid copy number and strength of constitutive expression; this would require no need for the modification of protein sequence/structure or binding affinities. In addition, a successful tuning strategy based on this approach may be portable to other TCS biosensors in future studies. Therefore, we set out to investigate whether varying the concentrations of AtoS HK, AtoC RR and Pato promoter would result in changes in final biosensor response, that agree with the predictions returned by sensitivity analysis of the AtoSC TCS model. Further work could explore the modification of other parameters identified as important for determining biosensor behaviour using sensitivity analysis (Figure 2C).

### Design and construction of an acetoacetate-inducible biosensor

As described above, the AtoSC TCS consists of the AtoS HK, AtoC RR and Pato promoter. Initially, a basic AtoSC biosensor was constructed by incorporating GFP expression under the control of the Pato promoter on a high-copy pUC based plasmid, designated ASAH0 (Figure 3A). The ASAH0 plasmid relies on host genomic expression of both AtoS and AtoC. The ASAH0 plasmid was initially characterised in three *atoSC*^+^ *E. coli* strains. As can be seen from Figure 3B and 3C, ASAH0 showed a robust increase in GFP expression with increasing acetoacetate concentration in all three strains. The fitted Hill parameters for each of these strains are given in SI Table S5. The response was similar in all three hosts, with NEB5*α* ASAH0 displaying higher fitted *f^max^* and *K*_1/2_ values. Furthermore, the ASAH0 plasmid showed a *K*_1/2_ commensurate with the concentrations of ketone bodies expected in human blood (sub-millimolar concentrations), indicating that they are responsive to acetoacetate at physiologically relevant levels[26]. However, ASAH0 was unresponsive to acetoacetate in *atoSC*^−^ (BW28878), *atoS*^−^ (JW2213) and *atoC*^−^ (JW2214) strains (SI Figure S2). This indicates that all components of the AtoSC TCS are required for the detection of acetoacetate, and that no other TCSs activate expression from the Pato promoter under these conditions.

**Figure 3:**
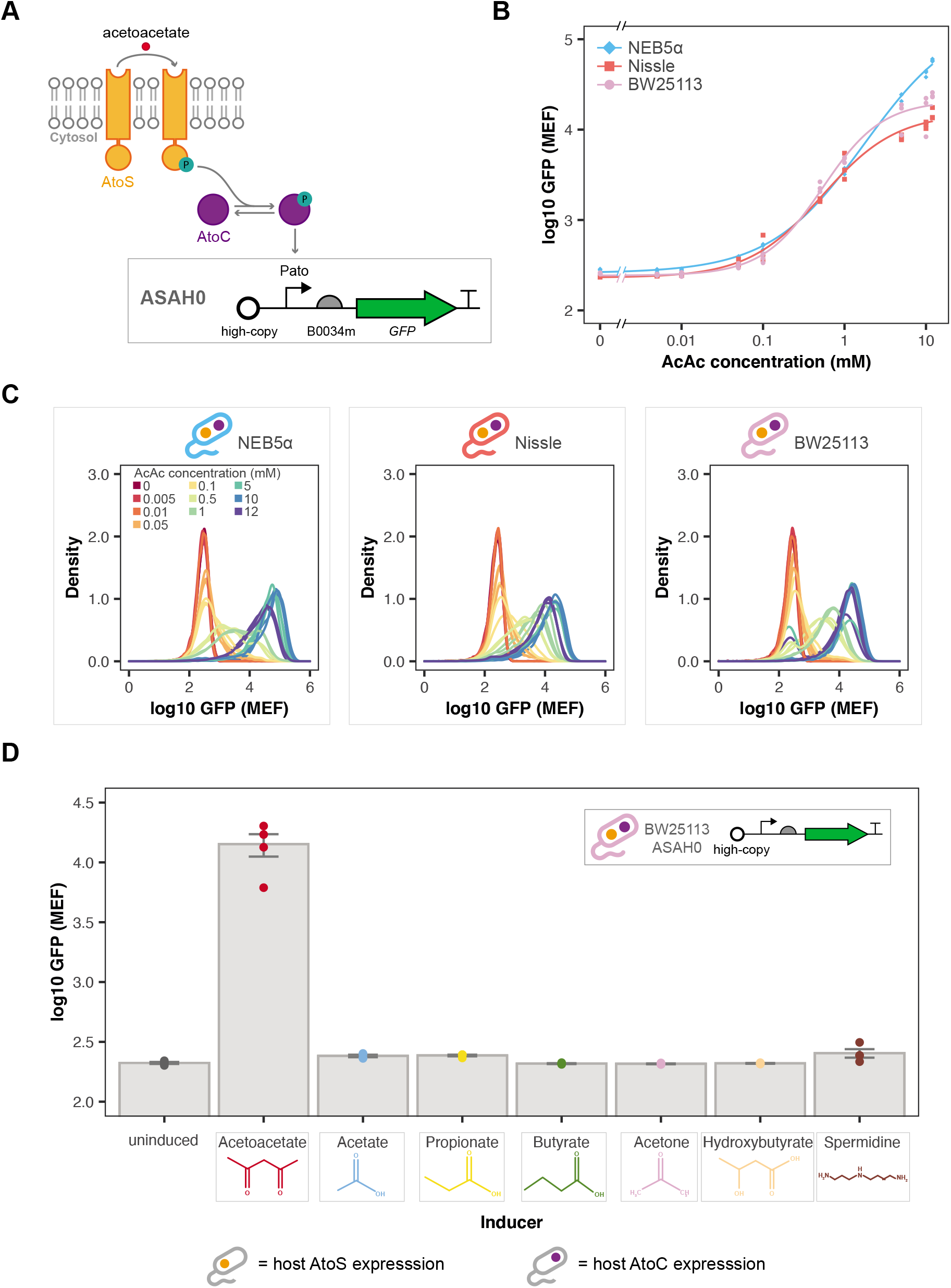
Characterisation of the acetoacetate-sensitive ASAH0 plasmid. (**A**) ASAH0 contains GFP under the control of the Pato promoter and relies on host expression of the AtoS HK and AtoC RR proteins. (**B**) Median GFP response of ASAH0 in three *atoSC*^+^ strains: NEB5*α*, BW25113 and Nissle 1917, exposed to acetoacetate. (**C**) Density plots of GFP fluorescence for ASAH0 in the three *atoSC*^+^ host strains (n = 3 biological repeats, data fit with Hill function). (**D**) GFP response of BW25113 ASAH0 to a range of alternative inducers in LB media, all at 20 mM (n = 4 biological repeats, points show medians and bars mean of medians ± SE, strain symbols denote host expression of *atoS/atoC*).

### Specificity of the acetoacetate-inducible biosensor

After demonstrating that the AtoSC TCS could be used to construct acetoacetate-inducible biosensors (Figure 3B), alternative inducers were examined to check the specificity of the ASAH0 circuit. Six alternative inducers were tested, including the SCFAs acetate, propionate and butyrate, the other ketone bodies, acetone and *#x03B2;*-hydroxybutyrate, and spermidine (which has previously been reported to act as an inducer for the AtoSC TCS)[43]. All were tested at a concentration of 20 mM, a concentration above that shown to give maximum acetoacetate induction. When characterised in LB media, acetoacetate produced the highest increase in GFP expression (Figure 3D), while the alternative inducers showed similar levels of GFP to the uninduced control. As 20 mM is a relatively high concentration, growth assays were measured to ensure the lack of GFP was not caused by toxicity of any of the inducers. All inducers produced no substantial changes in growth compared to the uninduced control, except for spermidine (SI Figure S3), which was seen to prevent cell growth. To further examine spermidine induction of BW25113 ASAH0, a full response curve was collected across a range of inducer concentrations (SI Figure S4). However, no increase in GFP was seen across this range of spermidine concentrations.

Characterisation was also performed for the BW25113 ASAH0 biosensor in several different media types (SI Figures S5 and S6). Firstly, the switching of BW25113 ASAH0 in minimal media with glycerol and glucose carbon sources was compared to that in LB media (SI Figure S5). The biosensor functioned across all three media types, although the fold change was greatly reduced with glucose as the carbon source. In addition, to determine whether the ASAH0 biosensor was capable of operating in more complex conditions, the biosensor was characterised in media supplemented with a range of mammalian culture media and HeLa/CHO-K1 conditioned samples (SI Figure S6). BW25113 ASAH0 was found to show induction when exposed to acetoacetate in cultures supplemented with 20% of these complex samples.

### Plasmid vs genome expression of AtoS/AtoC components

The ASAH0 plasmid relies on host expression of both the AtoS HK and AtoC RR proteins. However, motivated by the results of the sensitivity analysis, we wished to create versions of the AtoSC biosensor where the concentrations of both proteins could be varied (through changes in plasmid copy number). ASAH2J06 was generated through addition of the *atoS/atoC* genes to the ASAH0 plasmid, under the control of a constitutive promoter (J23106 of the Anderson promoter library: parts.igem.org); in a manner analogous to a previously reported nitrate biosensor that incorporated the *narX/narL* genes[11]. All other plasmid components were kept the same. The layout of this biosensor is given in Figure 4A. In contrast to ASAH0, the ASAH2J06 biosensor produced an acetoacetate-inducible change in GFP expression even when host expression of both the *atoS* and the *atoC* genes were knocked-out (SI Figure S7). In order to explore the effect of plasmid vs host gene expression in more detail the responses of ASAH0 in BW25113 (only host expression of AtoS/AtoC) and ASAH2J06 in BW28878 (only plasmid expression of AtoS/AtoC) were directly compared (Figure 4B). BW25113 ASAH0 displayed a greater *f^max^*, dynamic range and lower *f^min^* than ASAH2J06. However, BW28878 ASAH2J06 showed a fitted *K*_1/2_ value an order of magnitude lower than the BW25113 ASAH0 biosensor (31.7 ± 3.78 vs 344 ± 42.1 *μ*M, respectively). The source of AtoS HK and AtoC RR expression were the only differences between these strains. However, these relatively simple changes produced a dramatic shift in the response curves of the BW25113 ASAH0 and BW28878 ASAH2J06 biosensors, supporting the predictions gained through sensitivity analysis of the AtoSC model.

**Figure 4:**
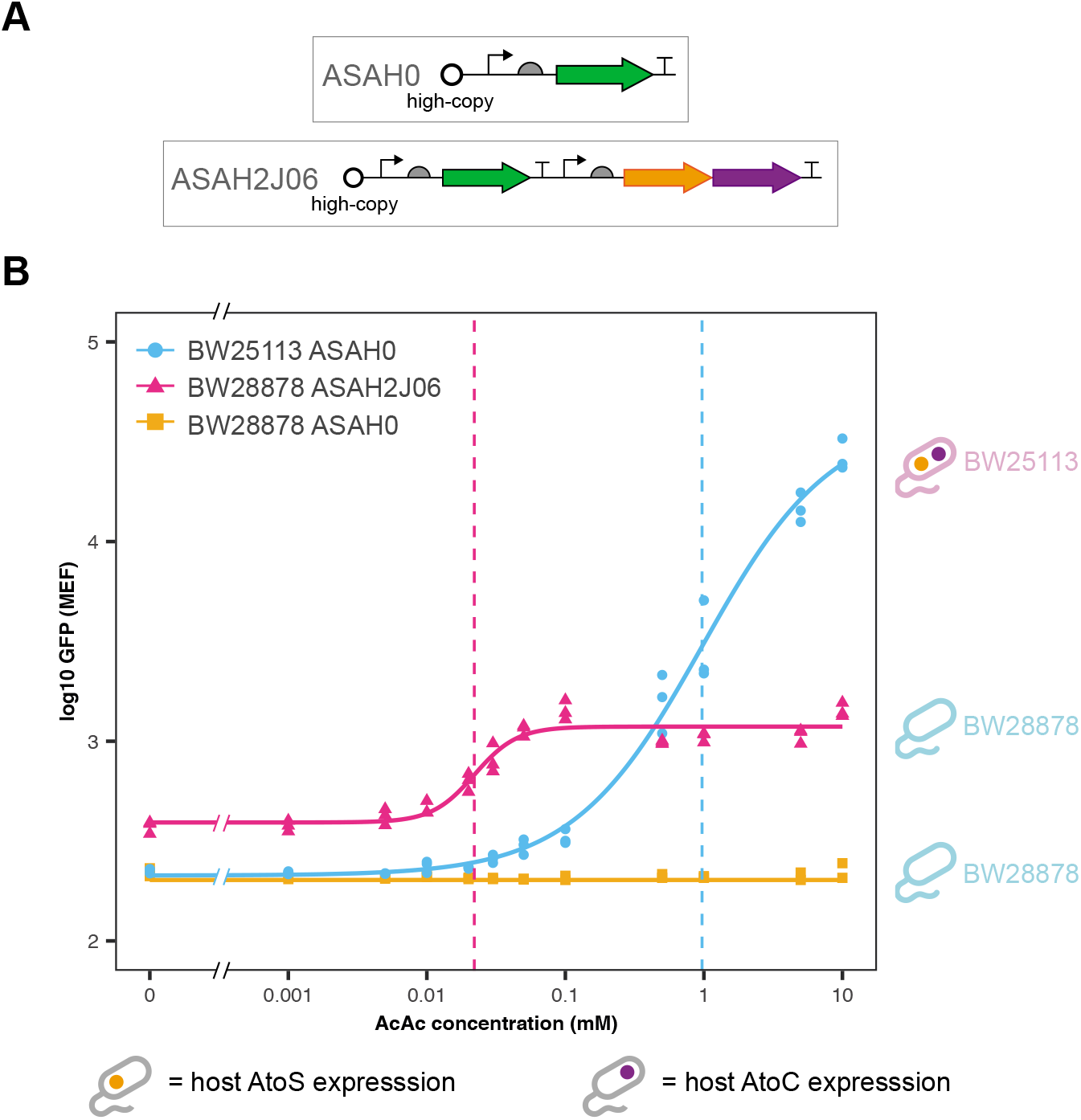
A comparison of the acetoacetate biosensor response between genomic and plasmid expression of the AtoS and AtoC genes. (**A**) Layouts of the ASAH0 and ASAH2J06 plasmids. ASAH2J06 contains constitutive expression of the AtoS HK and AtoC RR proteins. (**B**) Median GFP fluorescence of the whole-cell biosensors exposed to acetoacetate (BW25113 ASAH0 and BW28878 ASAH2J06: n = 3 biological repeats, data fit with Hill function, dashed lines indicate the fitted *K*_1/2_ values, BW28878 ASAH0: n = 2 biological repeats, line indicates lowest measured GFP median, cell symbols indicate the *atoS* and *atoC* presence status in the host chassis genome for each of the three whole-cell biosensor strains).

### Engineering modified biosensor responses by varying AtoS, AtoC and Pato concentrations

As the BW25113 ASAH0 and BW28878 ASAH2J06 biosensors displayed such different responses to acetoacetate induction we wished to explore if further changes to the concentrations of AtoS, AtoC and the Pato promoter would result in other behaviours. To this end, a range of further AtoSC plasmids were produced. Low copy versions of both the ASAH0 and ASAH2J06 plasmids were created by replacing the origin of replications with a low copy SC101 origin. These low copy plasmids were designated ASAL0 and ASAL2J06, respectively. Furthermore, the *atoC* genes were removed from the ASAH2J06 and ASAL2J06 plasmids to produce the ASAH1J06 and ASAL1J06 plasmids, which have plasmid expression of the AtoS HK but rely on host expression of the AtoC RR. The response curves for all six of these biosensors are given in Figure 5.

**Figure 5:**
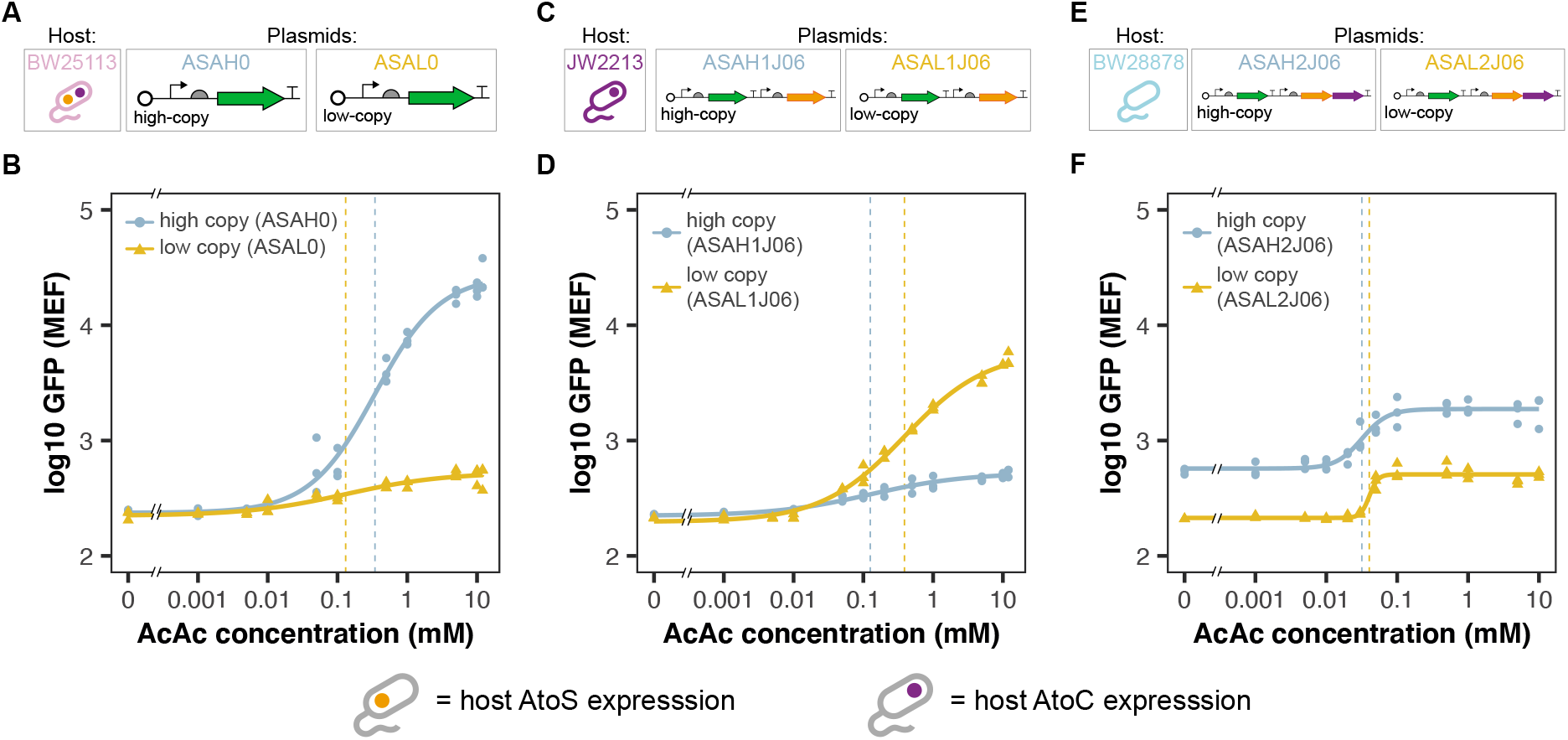
Manipulating whole-cell biosensor response by changing the copy number of components of the AtoSC two-component system. (**A**) Plasmid layouts of ASAH0 and ASAL0. (**B**) Median GFP response of ASAH0 and ASAL0 in the *atoSC*^+^ BW25113 host strain. (**C**) Plasmid layouts of ASAH1J06 and ASAL1J06. (**D**) Median GFP response of ASAH1J06 and ASAL1J06 in the *atoS*^−^ JW2213 host strain. (**E**) Plasmid layouts of ASAH2J06 and ASAH2J06. (**F**) Median GFP response of ASAH2J06 and ASAL2J06 in the *atoSC*^−^ BW28878 host strain (n = 3 biological repeats, data fit with Hill function, dashed lines indicate fitted K_1/2_ values).

Increasing the copy number of the plasmid containing only the Pato promoter produced a large increase in the *f^max^* of the final GFP response, while having no noticeable effect on *f^min^* (Figure 5B). Increasing the copy number of the plasmid containing both Pato and the AtoS HK (ASAH1J06/ASAL1J06) reversed this trend, with the lower copy number displaying a higher *f^max^*. Again little difference was seen in *f^min^* (Figure 5D). Finally, increasing the copy number of the plasmid containing Pato, the AtoS HK and AtoC RR (ASAH2J06/ASAL2J06) resulted in a vertical shift of the GFP response, increasing both *f^min^* and *f^max^* (Figure 5F). All of the fitted parameters to these response curves are given in SI Table S5. Of all the *ato* knockout (BW25113, BW28878, JW2213 and JW2214) whole-cell biosensor strains constructed, BW25113 ASAH0 displayed the largest dynamic range (114), whereas BW28878 ASAH2J06 was found to have the lowest *K*_1/2_ (31.7±3.78 *μ*M) followed by the low-copy BW28878 ASAL2J06 (40.1±2.28 *μ*M). From these response curves it is clear that introducing variations in the concentrations of AtoS HK, AtoC RR and Pato promoter, through varying plasmid vs host genome expression and plasmid copy number, can produce dramatically different response curves and associated behaviours, as qualitatively predicted by sensitivity analysis of the AtoSC model.

## Discussion

### Insights from sensitivity analysis

The AtoSC TCS is a complex system with a large number of different species and parameters involved in its functioning. This presents a challenging task when trying to decide which parameters to modify during circuit engineering. Morris sensitivity analysis was performed to try and identify the parameters which might have the greatest influence on biosensor behaviour, and therefore the largest scope for effective engineering of the final biosensor response. When ranking the Morris sensitivity analysis all parameters involving the reporter elements (e.g. GFP maturation/degradation) were removed from the results. Biosensors can be built with a range of reporters and outputs[44]; therefore, we wished to focus on the parameters influencing the TCS, as these features may have wider value for engineering other TCS based biosensors. From the final rankings, it can be seen that the concentration of AtoC RR (‘C’), the concentration of AtoS HK (‘S’), and Pato promoter concentration (‘pato’), were predicted to have an effect on a range of different aspects of biosensor behaviour (Figure 2C). These results provided motivation for the whole-cell biosensor designs characterised within Figures 4 and 5, as changing the availability of these three main components was hypothesised to have a large effect on biosensor behaviour. It should be noted that other parameters were also predicted to have an effect on biosensor behaviour (e.g. the binding rate of the phosphorylated AtoC RR to the Pato promoter). It may be possible to use these so far unexplored parameters as a starting point for further attempts to engineer acetoacetate biosensors with different responses in the future. In addition, further work could include the fitting of TCS parameters from time-course data, thereby improving the predictive capability of the model.

### Using the AtoSC TCS to create acetoacetate-inducible biosensors

The ASAH0 plasmid is the simplest design for an acetoacetate sensitive biosensor that can be created from the AtoSC TCS, requiring host expression of the AtoS HK and AtoC RR. ASAH0 was initially tested in three *E. coli* strains: NEB5*α* (a commercial cloning strain), BW25113 (the parent strain of the Keio knockout collection[45]) and Nissle 1917 (a commonly used microbiome engineering strain[46]). Although the ASAH0 design is simple and was found to be acetoacetate-inducible in all three strains, it provides fewer components that can be modified to tune biosensor performance. The aim of the ASAH2J06 design was to provide a self contained plasmid that would (a), not rely on host expression of AtoS HK/AtoC RR, and (b), provide a greater potential design space from which to produce the desired biosensor behaviour. Upon construction both ASAH0 and ASAH2J06 were found to be acetoacetate-inducible (Figure 4).

As mentioned previously, the typical concentrations of ketone bodies within a healthy adult are sub-millimolar. It has been reported that these may increase to approximately 1.0 mM during hyperketonemia, above 3.0 mM during ketoacidosis and reach levels in excess of 20 mM during cases of uncontrolled diabetes[27, 47]. The original form of the whole-cell biosensor, BW25113 ASAH0, was able to sense and respond to acetoacetate concentrations within this physiologically relevant range, and displayed a *K*_1/2_ lower than the detection limit of the commercially available Ketostix, which rely on a sodium nitroprusside reaction to detect levels of acetoacetate in a semi-quantitative manner (*K*_1/2_ of 344±42.1 *μ*M vs detection limit of ~500 *μ*M, respectively). These are desirable characteristics if the biosensor is to be used for diagnostic applications. However, it should be noted that the relative levels of the different ketone bodies may change across different diseased states. For example, in a healthy adult the ratio of *β*-hydroxybutyrate to acetoacetate commonly lies between 1:1 to 3:1; however, during cases of acute diabetic ketoacidosis this ratio can vary greatly due to impaired interconversion of *β*-hydroxybutyrate to acetoacetate[47]. Therefore, the actual proportion of acetoactetate may vary depending on the disease state. This means that more sensitive biosensors (i.e. biosensors with a lower *K*_1/2_) may be required for certain diagnostic applications. As can be seen from Figure 4, the modified BW28878 ASAH2J06 biosensor showed a substantially lower *K*_1/2_ than the original BW25113 ASAH0 (31.7±3.78 vs 344±42.1 *μ*M, respectively). As expected, BW28878 ASAH0 showed no response to acetoacetate induction (Figure 4 and SI Figure S2), as the AtoS HK and AtoC RR are not expressed (either from the genome or plasmid) within this strain. These results suggest that replacing host genome with plasmid expression could offer a starting point for attempts to reduce the *K*_1/2_ of other TCS based whole-cell biosensors.

The BW25113 ASAH0 biosensor was found to be specific to acetoacetate when tested against a range of alternative inducers. As the AtoSC TCS and the genes of the *atoDAEB* operon are known to play a role in SCFA metabolism the SCFAs acetate, propionate and butyrate were tested as alternative inducers. The data collected here supports previous results that have reported SCFAs, particularly butyrate, can not induce the AtoSC TCS[48, 49]. In addition, the two other ketone bodies, acetone and *β*-hydroxybutyrate were tested for induction; neither were found to induce the BW25113 ASAH0 biosensor. As discussed previously, spermidine has been reported elsewhere as an alternative inducer of the AtoSC TCS[43]. Characterisation of BW25113 ASAH0 did not show an increase in GFP response when exposed to spermidine (Figure 3D and SI Figure S4). However, it is possible that the spermidine concentrations needed to trigger GFP production from this plasmid may be above those that become toxic to the cells. Future work could explore whether spermidine induction of the BW25113 ASAH0 biosensor can be seen in other culture conditions. Biosensors may behave differently when exposed to different culture conditions. In order to further explore the switching response of BW25113 ASAH0, the biosensor was characterised in a range of more complex media components (SI Figures S5 and S6). BW25113 ASAH0 displayed an increase in GFP upon exposure to acetoacetate in all the conditions tested; however, the fold change of the switching was found to be much lower when grown in M9 media with glucose as the carbon source (SI Figure S5). Previous work has shown that the *atoD* gene (present at the beginning of the *atoDAEB* operon) is a target of the Hfq-binding small RNA, Spot 42, which suppresses numerous metabolic genes in the presence of a preferred carbon source[50]. This is a likely mechanism for the repression seen with the BW25113 ASAH0 strain. Future work could explore methods of reducing this catabolic repression within the engineered whole-cell biosensor strains.

## Conclusions

Our study has demonstrated that the AtoSC TCS can be used to construct acetoacetate-inducible biosensors. Characterisation of the BW25113 ASAH0 whole-cell biosensor showed that the biosensor is specific to acetoacetate, with no GFP response recorded when exposed to a range of alternative inducer molecules. Sensitivity analysis performed on a model of the AtoSC TCS predicted that varying AtoS HK, AtoC RR and Pato promoter presence would have a large effect on the final biosensor response. Subsequently, we constructed a range of AtoSC whole-cell biosensors with varying expression of AtoS HK, AtoC RR and presence of the Pato promoter. These whole-cell biosensors were found to exhibit different response curves when exposed to acetoacetate, supporting the insights gained through sensitivity analysis. In the future, these AtoSC whole-cell biosensors may be incorporated into a range of other biomedical, bioproduction or agricultural applications, where there is a need to monitor the levels of the acetoacetate ketone body. The design methods used here may also be applied to the construction of other TCS biosensors, where there is a mismatch between the desired and observed biosensor behaviour.

## Notes

### Competing Interest Statement

The authors have declared no competing interest.

### Summary of Updates

A fully updated manuscript describing new constructs and modelling.

