## Supplementary Information for "Engineered acetoacetate-inducible whole-cell biosensors based on the AtoSC two-component system"

### 1 Extended Materials and Methods

#### 1.1 Strains & Primers

Table S1 details the primers used to construct the Ato WCBs in this work. Table S2 details the host strains and plasmids used within this work.

#### 1.2 Ato biosensor plasmid construction

All plasmids were constructed using CIDAR MoClo and general cloning techniques. A MoClo transcription unit containing the Pato promoter and GFP was constructed from the LD1 pAto DVA_EB, DVK_EF, B0034m_BC, E0040m_CD and B0015_DF (CIDAR MoClo kit) DNA parts. This unit was designated pAto + GFP. The ASAH0 plasmid was constructed by adding the pAto + GFP transcriptional unit, into the DVA_EF MoClo backbone plasmid. The low-copy ASAL0 plasmid was made by amplifying the SC101 origin from the DVS471_AF plasmid, using the SC101_fragment.F and SC101_fragment.R primers (SI Table S1). ASAH0 was amplified using the SC101_vector.F and SC101_vector.R primers. The fragments were then combined using HiFi DNA assembly (New England Biolabs, following the manufacturers protocol). The AtoS/AtoC fragment was amplified from *E. coli* genomic DNA using the LD9 and LD10 primers and inserted into the DVA_CD MoClo backbone to produce AtoSC_CD. A transcription unit was then created using AtoSC_CD, DVK_FG, J23106_FB,

Table S1: Oligonucleotides used within this study. Lowercase letters indicate the annealing sequence. Underlined letters show addition of restriction enzyme cleavage sites.

| **Oligonucleotide** | **Sequence** (5' to 3') |
|---|---|
| SC101_fragment.F | TGGCGTTcttttccgctgcataaccctgc |
| SC101_fragment.R | AAAGGATCTTCgagttatacacagggctgggatctatt |
| SC101_vector.F | GTGTATAACTCgaagatcctttgatcttttctacggggt |
| SC101_vector.R | CGGAAAAGaacgccagcaacgcggc |
| LD9 | GGCGGTCTCA<u>AATG</u>atgcattatatgaagtggatttatccacgcc |
| LD10 | GGCGGTCTCT<u>ACCT</u>TCAttatacatccgccggatc |
| LD89 | GGCGGTCTCT<u>ACCT</u>catacagtctgatttccctgcgg |

Table S2: Bacterial strains and plasmids used within this study.

| | **Description** | **Source** |
|---|---|---|
| **Host strain** | | |
| *E. coli* NEB5$\alpha$ | Cloning strain | New England BioLabs |
| *E. coli* BW25113 | Keio collection parent strain | Keio collection |
| *E. coli* JW2213 | Keio collection *atoS* knockout | Keio collection |
| *E. coli* JW2214 | Keio collection *atoC* knockout | Keio collection |
| *E. coli* BW28878 | *atoS* and *atoC* double-knockout | Prof. Kyriakidis, University of Thessaloniki |
| *E. coli* Nissle 1917 | Commensal *E. coli* strain | Prof. Ian Henderson, University of Birmingham |
| **Plasmid** | | |
| LD1 pAto | DVA_EB CIDAR MoClo backbone harbouring the Pato promoter | This work |
| pAto + GFP | As LD1 pAto, with GFP | This work |
| DVS471_AF | Low-copy SC101 origin plasmid | Lab stock |
| AtoSC_CD | DVA_CD CIDAR MoClo backbone harbouring the *atoSC* genes | This work |
| ASAH0 | High-copy pUC origin, Pato promoter controlled GFP expression, amp$^R$ | This work |
| ASAL0 | Low-copy SC101 origin, Pato promoter controlled GFP expression, amp$^R$ | This work |
| ASAH1J06 | As ASAH0, with constitutive J23106 promoter driving AtoS expression | This work |
| ASAL1J06 | As ASAL0, with constitutive J23106 promoter driving AtoS expression | This work |
| ASAH2J06 | As ASAH0, with constitutive J23106 promoter driving AtoS and AtoC expression | This work |
| ASAL2J06 | As ASAL0, with constitutive J23106 promoter driving AtoS and AtoC expression | This work |

B0034m_BC and B0015_DG. This transcription unit was designated AtoSC unit. The ASAH2J06 plasmid was made by combining the AtoSC and pAto + GFP units in the DVA_EG backbone. As with ASAL0, the low-copy ASAL2J06 plasmid was made by amplifying the SC101 origin from the DVS471_AF plasmid, using the SC101_fragment.F and SC101_fragment.R primers (SI Table S1). ASAH2J06 was amplified using the SC101_vector.F and SC101_vector.R primers. The two fragments were then combined using HiFi DNA assembly. The AtoS fragment was amplified from ASAH2J06 using the LD9 and LD89 primers (SI Table S1). This was then combined with DVK_FG, J23106_FB, B0034m_BC and B0015_DG, to produce the AtoS unit. Again, the low-copy ASAL1J06 plasmid was made by amplifying the SC101 origin from the DVS471_AF plasmid, using the SC101_fragment.F and SC101_fragment.R primers. ASAH1J06 was amplified using the SC101_vector.F and SC101_vector.R primers. The two fragments were then combined using HiFi DNA assembly. Maps of the biosensor plasmids constructed are given in Figure S9.

##### 1.3 Complex media assays

Induction assays of the BW25113 ASAH0 plasmid were performed in a variety of media types, to explore performance of the biosensor in different operating conditions. Minimal M9 media (10 ml: 5x M9 salts, 1 ml: 10% casamino acids, 100 $\mu$L: 1M $MgSO_4$, 5 $\mu$L: 1M $CaCl_2$, per 50 ml of media) was prepared with either glucose (400 $\mu$L: 50% glucose) or glycerol (250 $\mu$L: 80% glycerol) as the carbon source. The induction assays were then performed following the method given in the main text, with LB media replaced with the desired culture media.

Surrogate Dulbecco's Modified Eagle Media (DMEM) without phenol red was made as follows: (100 ml: 0.83 g DMEM powder (SIGMA, D5030-1L),25 mM D-glucose, 10 m HEPES pH7.4, 2 mM Glutamax (Gibco 35050038)). This was then filter sterilised before use and stored at 4°C. Pre-made DMEM Media containing Phenol red (DMEM High glucose, HEPES, Glutamax, Gibco 32430-027) was collected before FBS (10%, Gibco, 10270) was added, and another sample collected. Pre-treated T75 flasks (Thermo 156499) containing 25 ml of this media were seeded with either HeLa cells (ECACC: 93021013 - below 10th passage) at $1x10^6$ cells or CHO-k1 cells (ECACC: 85051005) at $0.5x10^5$ cells with L-Proline addition (Sigma, p5607) and incubated for 48h. After growth at 37°C, 5% $CO_2$ in a standard stack incubator, media was collected from flasks and centrifuged for 10 minutes at 5000 g to pellet cell material. A 10 ml sample of media supernatant was then collected and stored at 4°C, prior to induction assays. Assays involving these complex media samples were performed based on a modified version of the protocol reported by Goers *et al.* (2017)[1]. Overnight cultures were diluted to an $OD_{700}$ of approximately 0.05 in M9 glycerol media, and 150 $\mu$L added to each well of a polypropylene 96 deep-well plate (Brand, Sigma Aldrich). The plate was then incubated for 2 hours at 37°C, with 350 rpm shaking, sealed with an autoclaved system Duetz lid. Each well was then supplemented with 40 $\mu$L of the desired complex media, before inducers added to the desired concentration, bringing the total volume in each well to 200 $\mu$L (giving a final media sample concentration of 20% as in Goers *et al.*[1]). The plate was once again sealed with an autoclaved system Duetz lid and the induced cultures incubated at 37°C, with 350 rpm shaking for 16 hours. Samples were taken as required for flow cytometry analysis.

All complex media induction assays were normalised against untransformed BW25113 host strain cultures grown in the corresponding media conditions. The control was run in triplicate and BW25113 ASAH0 data normalised by subtracting the mean of three median control measurements. The standard error for these measurements was calculated as the propagated standard error, following the variance formula:

$$SE_p{}^2 = SE_s{}^2 + SE_c{}^2 \tag{1}$$

where $SE_p$ is the propagated standard error, $SE_s$ the standard error of the sample and $SE_c$ the standard error of the control.

#### 2 Development of the AtoSC TCS model.

A chemical reaction network model was constructed to try and capture the behaviour of the AtoSC whole-cell biosensors. This model was originally adapted from a previously published TCS model[2], through the addition of reactions describing the expression of GFP from the biosensor plasmid. This model described the well-characterised EnvZ/OmpR TCS; chosen as this TCS produces a graded response (similar to AtoSC) and is a prototypical TCS that does not require auxiliary proteins[2]. An overview of the model parameters and species that were used to describe the AtoSC TCS behaviour is given in Figure 2, of the main text.

Firstly, AtoS can be autophosphorylated in the presence of acetoacetate, in the reverse of this reaction it can also be dephosphorylated

$$\text{AtoS} \underset{k_{ad}}{\overset{k_{ap}}{\rightleftharpoons}} \text{AtoS}^* \tag{2}$$

Within this model the phosphorylation rate, $k_{ap}$, is determined by the concentration of acetoacetate inducer; therefore, this rate was varied to simulate changing levels of acetoacetate inducer (as in the original study, from which this model was adapted)[2]. Following the autophosphorylation of AtoS, the phosphoryl group can be transferred between the AtoC response regulator. Firstly, AtoC binds to phosphorylated AtoS

$$\text{AtoS}^* + \text{AtoC} \underset{k_{d1}}{\overset{k_{b1}}{\rightleftharpoons}} \text{AtoS}^*.\text{AtoC} \tag{3}$$

The phosphoryl group is then transferred, followed by disassociation of the AtoS-AtoC complex. Alongside transferring the phosphoryl group to the RR, some HKs have been shown to dephosphorylate their cognate RR in the absence of their inducer. Previous reports have implied that the AtoC RR may interact with the HK of another TCS system *in vitro*[3]. Although a later study did not find evidence of this interaction *in vivo*[4]. As a simplifying assumption, we disregard the alternative dephosphorylation pathway and only include dephosphorylation of AtoC* by unphosphorylated AtoS

$$\text{AtoS}^*.\text{AtoC} \underset{k_{ph}}{\overset{k_{pt}}{\rightleftharpoons}} \text{AtoS}.\text{AtoC}^* \tag{4}$$

$$\text{AtoS}.\text{AtoC}^* \underset{k_{b2}}{\overset{k_{d2}}{\rightleftharpoons}} \text{AtoS} + \text{AtoC}^* \tag{5}$$

In addition, it has been shown that it is possible for HKs to bind to their RR even in the absence of a phosphoryl group. This leads to the formation of a 'deadend' complex, which effectively sequesters RR from the system

$$\text{AtoS} + \text{AtoC} \underset{k_{d3}}{\overset{k_{b3}}{\rightleftharpoons}} \text{AtoS}.\text{AtoC} \tag{6}$$

The phosphorylated AtoC can bind to the $\text{p}_{\text{ato}}$ promoter to induce transcription

$$\text{AtoC}^* + \text{p}_{\text{ato}} \underset{k_{unb}}{\overset{k_{bnd}}{\rightleftharpoons}} \text{AtoC}^*.\text{p}_{\text{ato}} \tag{7}$$

Finally, we include a set of simplified transcription/translation reactions which describe the production of GFP from the $\text{p}_{\text{ato}}$ promoter. Leaky transcription from the uninduced $\text{p}_{\text{ato}}$ promoter is included to capture basal expression levels

$$\mathrm{AtoC^*.p_{ato}} \overset{k_{pmgexp}}{\rightarrow} \mathrm{mGFP + AtoC^*.p_{ato}} \tag{8}$$

$$\mathrm{p_{ato}} \overset{k_{lmgexp}}{\rightarrow} \mathrm{mGFP + p_{ato}} \tag{9}$$

$$\mathrm{mGFP} \overset{k_{gexp}}{\rightarrow} \mathrm{mGFP + GFP} \tag{10}$$

$$\mathrm{mGFP} \overset{k_{mdeg}}{\rightarrow} \emptyset \tag{11}$$

$$\mathrm{GFP} \overset{k_{deg}}{\rightarrow} \emptyset \tag{12}$$

The above chemical reaction network was compiled into a set of ordinary differential equations, assuming mass-action kinetics. The results of sensitivity analysis for this model are shown in Figure S1.

#### Additional Ato WCB characterisation

Table S4 gives the fitted Hill parameters for each of the response curves shown in Figures 3 - 5 of the main text.

Figure S2 shows characterisation of the ASAH0 plasmid in the BW25113 ($atoSC^+$), JW2213 ($atoS^-$), JW2214 ($atoC^-$) and BW28878 ($atoSC^-$) strains. GFP induction was only seen in the BW25113 host strain. Figure S3 shows growth curves of the BW25113 ASAH0 whole-cell biosensor exposed to various inducers at 20 mM. No substantial changes in growth were seen, except for with spermidine which prevented growth of the biosensor strain. Figure S4 shows a comparison of induction curves of the BW25113 ASAH0 whole-cell biosensor exposed to acetoacetate and spermidine. No induction of GFP was seen across the range of spermidine concentrations tested.

Figure S7 shows characterisation of the ASAH2J06 plasmid in the BW25113 ($atoSC^+$), JW2213 ($atoS^-$), JW2214 ($atoC^-$) and BW28878 ($atoSC^-$) strains. GFP induction was seen in all host 'Ato' strains. Figure S8 gives concentration assays of the untransformed host strains; no change in GFP fluorescence on exposure to acetoacetate was seen with any of the four host strains.

Figure S5 shows normalised induction of the BW25113 ASAH0 whole-cell biosensor in three different media conditions: LB media and minimal M9 media with either glycerol or carbon sources. The annotated numbers give the fold change of the biosensor in each media. GFP induction was seen in all three medias, although was repressed in M9 with glucose. Figure S6 shows normalised induction of the BW25133 ASAH0 whole-cell biosensor in a range of additional complex medias. GFP induction was seen in all medias with exposure to acetoacetate. The largest fold change was observed in DMEM recording media, and the smallest in DMEM media supplemented with phenol red and 10% FBS.

Table S3: Morris sensitivity analysis parameter bounds. HK = histidine kinase (AtoS), RR = response regulator (AtoC), * denotes phosphorylation. Deliberately large ranges were selected for parameter bounds to try and capture the full range of possible biosensor responses.

| **Parameter** | **Description** | **Bounds** | **Ref** |
|---|---|---|---|
| | **Two-component system** | | |
| $k_{ap}$ | HK autophosphorylation | $10^{-9}$ - 10 $s^{-1}$ | |
| $k_{ad}$ | HK* autodephosphorylation | 0.0001 - 0.01 $s^{-1}$ | [2] |
| $k_{pt}$ | Phosphorylation of RR by HK* | 0.15 - 15 $s^{-1}$ | [2] |
| $k_{ph}$ | Dephosphorylation of RR* by HK | 0.005 - 0.5 $s^{-1}$ | [2] |
| $k_{b1}$ | Binding of HK* to RR | 0.05 - 5 $\mu M^{-1}$ $s^{-1}$ | [2] |
| $k_{d1}$ | Dissociation of bound HK* and RR | 0.05 - 5 $s^{-1}$ | [2] |
| $k_{b2}$ | Binding of HK and RR* | 0.05 - 5 $\mu M^{-1}$ $s^{-1}$ | [2] |
| $k_{d2}$ | Dissociation of bound HK and RR* | 0.05 - 5 $s^{-1}$ | [2] |
| $k_{b3}$ | Binding of HK and RR | 0.05 - 5 $\mu M^{-1}$ $s^{-1}$ | [2] |
| $k_{d3}$ | Dissociation of HK and RR | 0.05 - 5 $s^{-1}$ | [2] |
| $k_{bnd}$ | Binding of RR* to Pato promoter | 0.02 - 2 $\mu M^{-1}$ $s^{-1}$ | [5] |
| $k_{unb}$ | Unbinding of RR* and Pato promoter | 0.0005 - 0.05 $s^{-1}$ | [5] |
| | **Transcription & translation** | | |
| $k_{pmgexp}$ | Transcription of mRNA from induced Pato | 0.004 - 0.4 $s^{-1}$ | [6] |
| $k_{lmgexp}$ | Transcription of mRNA from uninduced Pato | 0.00004 - 0.004 $s^{-1}$ | [6] |
| $k_{gexp}$ | Translation of GFP from mRNA | 0.04 $s^{-1}$ | [6] |
| $k_{mgdeg}$ | mGFP degradation | 0.002 $s^{-1}$ | [6] |
| $k_{gdeg}$ | GFP degradation | 0.0004 $s^{-1}$ | [6] |
| | **Species concentrations** | | |
| $S_{tot}$ | *atoS* concentration | 0.001 - 0.1 $\mu$M | [6] |
| $C_{tot}$ | *atoC* concentration | 0.01 - 1 $\mu$M | [6] |
| $pato_{tot}$ | Total Pato promoter concentration | 0.001 - 0.1 $\mu$M | 1-100 per cell |

Table S4: Fitted parameter values to Hill functions for the GFP induction curves of the Ato WCBs (fitted values $\pm$ SE, given to 3 s.f.).

| **Host** | **Plasmid** | $f^{min}$ (MEF) | $f^{max}$ (MEF) | $K_{1/2}$ ($\mu$M) | $n$ | dynamic range |
|---|---|---|---|---|---|---|
| NEB$\alpha$ | ASAH0 | 259±1.04 | 190000±1.26 | 1760±206 | 0.744±0.0392 | 732 |
| EcN | ASAH0 | 230±1.07 | 15200±1.13 | 535±56.3 | 0.954±0.0965 | 64.8 |
| BW25113 | ASAH0 | 236±1.09 | 27100±1.14 | 344±42.1 | 0.928±0.0945 | 114 |
| BW25113 | ASAL0 | 223±1.05 | 528±1.08 | 131±61.6 | 0.614±0.164 | 1.37 |
| JW2213 | ASAH1J06 | 223±1.04 | 524±1.05 | 126±36.1 | 0.634±0.117 | 1.35 |
| JW2213 | ASAL1J06 | 198±1.04 | 5950±1.09 | 388±39.3 | 0.730±0.0505 | 29.1 |
| BW28878 | ASAH2J06 | 572±1.06 | 1880±1.05 | 31.7±3.78 | 2.38±0.636 | 2.28 |
| BW28878 | ASAL2J06 | 214±1.02 | 509±1.02 | 40.4±2.28 | 7.11±1.60 | 1.38 |

[6] Ron Milo, Paul Jorgensen, Uri Moran, Griffin Weber, and Michael Springer. BioNumbersthe database of key numbers in molecular and cell biology. *Nucleic Acids Research*, 38(suppl_1):D750–D753, jan 2010.

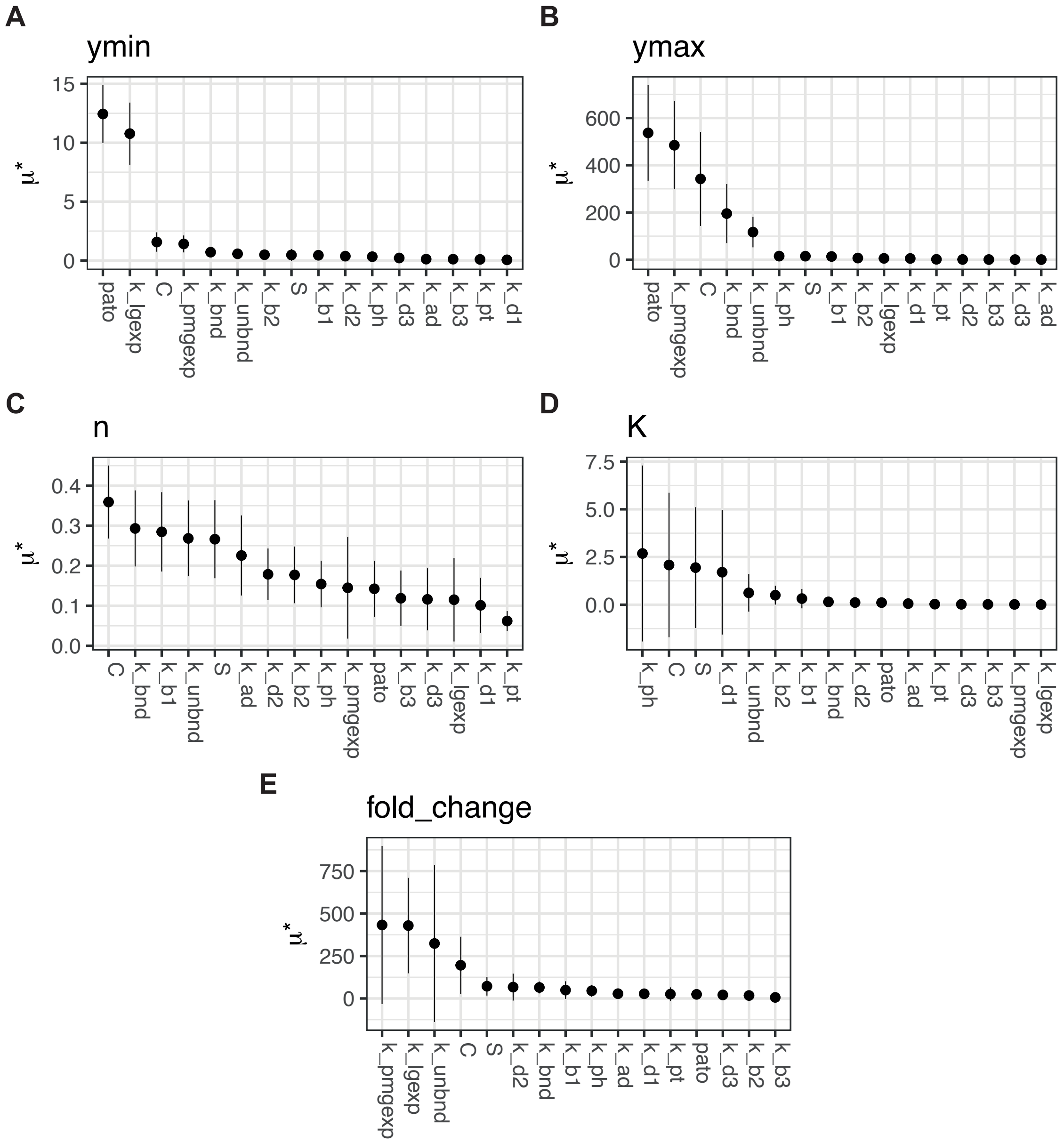


Figure S1: Sensitivity analysis rankings of the model parameters for different aspects of biosensor behaviour: (**A**) $f^{min}$/ymin, (**B**) $f^{max}$/ymax, (**C**) n, (**D**) $K_{1/2}$/K and (**E**) fold change. Rankings given as $\mu^*$ values $\pm$ 95% confidence intervals, a higher $\mu^*$ value predicts a greater impact on biosensor behaviour.

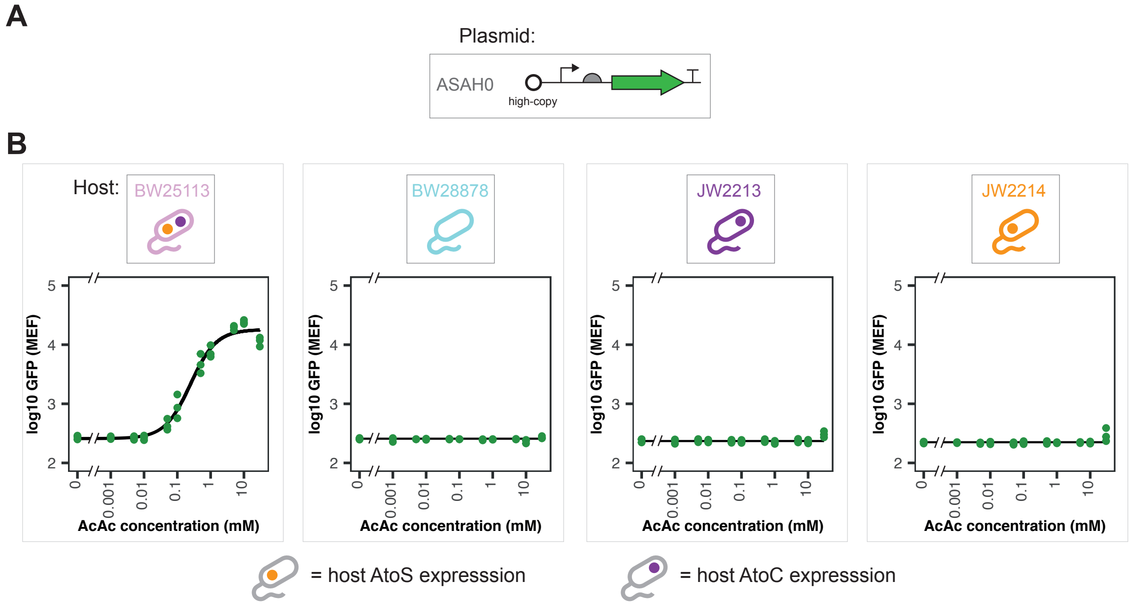


Figure S2: The performance of the ASAH0 biosensor plasmid in the four ‘Ato’ knockout strains. (**A**) Plasmid layout of ASAH0. (**B**) Median GFP fluorescence of the ASAH0 plasmid in each of the given host chassis, at varying acetoacetate concentrations. (n = 3 biological repeats, data fit with Hill function)

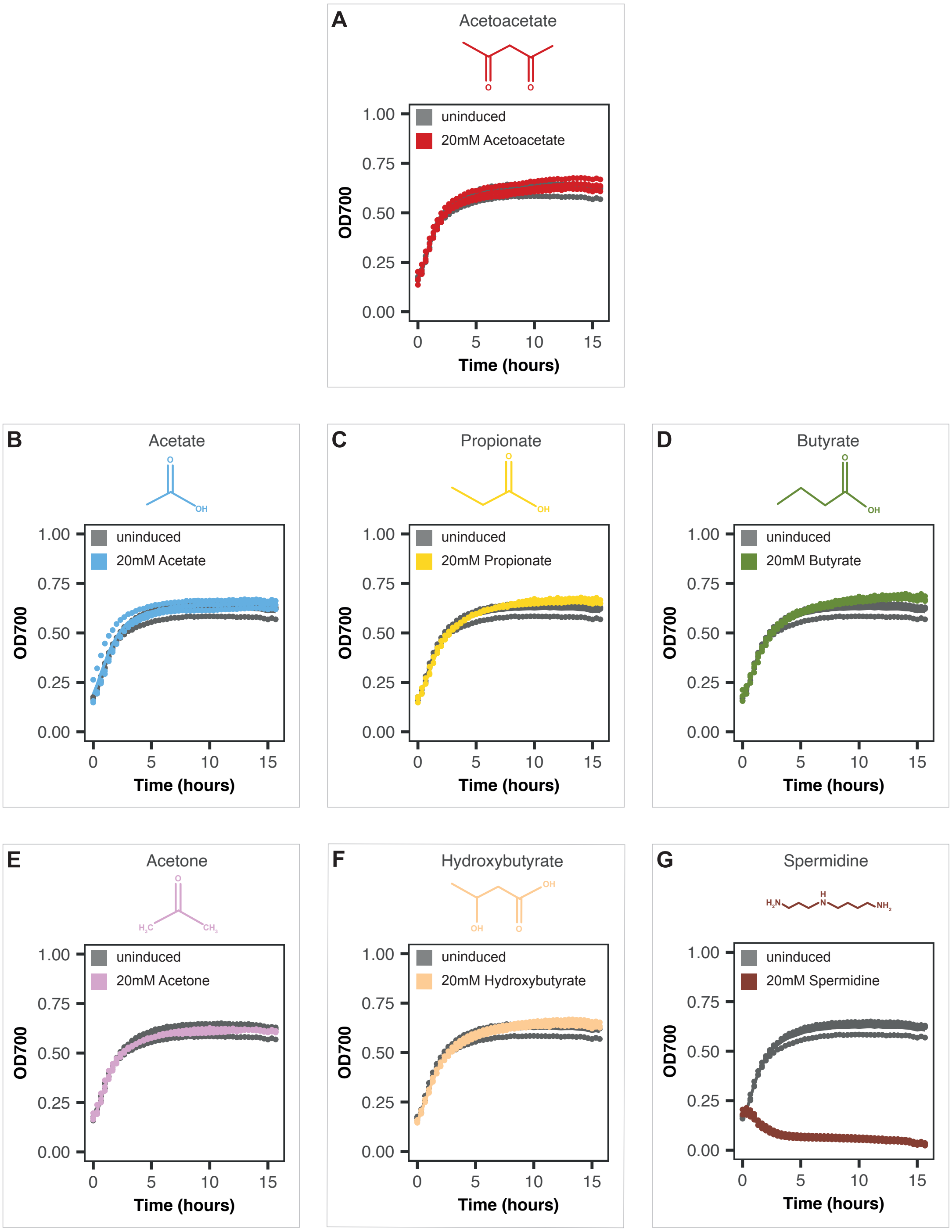


Figure S3: Growth curves of BW25113 ASAH0 exposed to a range of alternative inducer molecules, all at 20 mM within LB media, compared to that of the biosensor strain in unsupplemented LB media. Inducers: (**A**) acetoacetate, (**B**) acetate, (**C**) propionate, (**D**) butyrate, (**E**) acetone, (**F**) hydroxybutyrate and (**G**) spermidine. (n = 4 biological repeats, solid lines give means and points individual repeats)

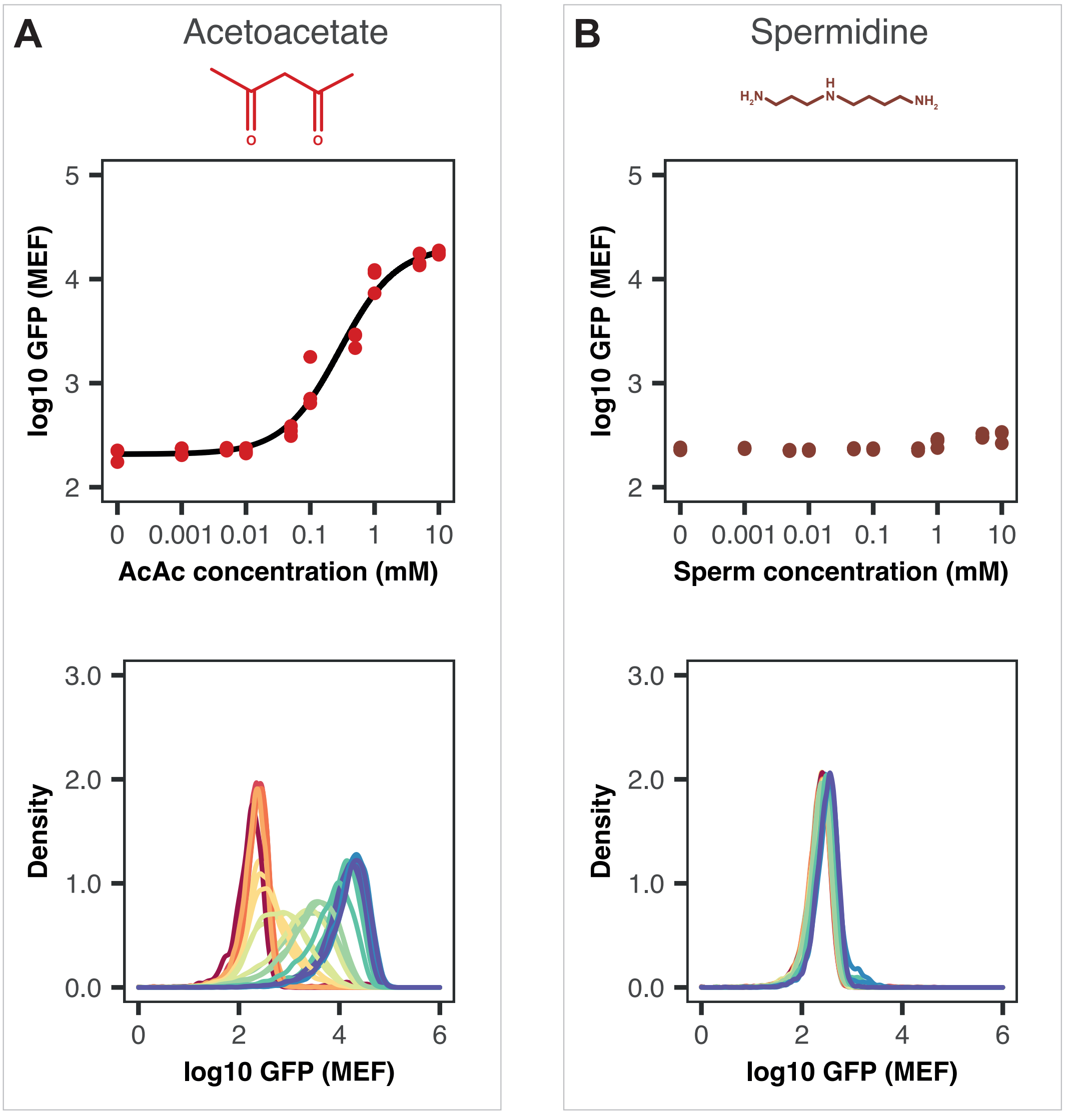


Figure S4: Concentration assays of the BW25113 ASAH0 whole-cell biosensor exposed to both (**A**) acetoacetate and (**B**) spermidine, within LB media. Top plots show median GFP fluorescence and bottom plots give density plots of GFP fluorescence. (n = 3 biological repeats, data fit to Hill equation)

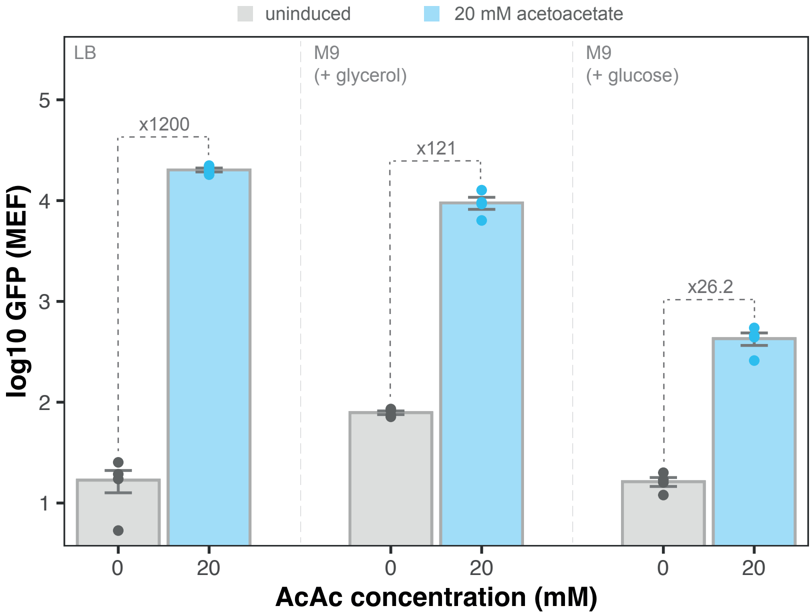


Figure S5: Normalised induction of the BW25113 ASAH0 whole-cell biosensor, in three medias: LB, M9 with glycerol and M9 with glucose carbon sources. The annotated numbers represent the fold change (calculated as mean normalised GFP at 20 mM induction divided by mean normalisedGFP at 0 mM induction). (n = 4 biological repeats, points show medians and bars means of medians ± SE)

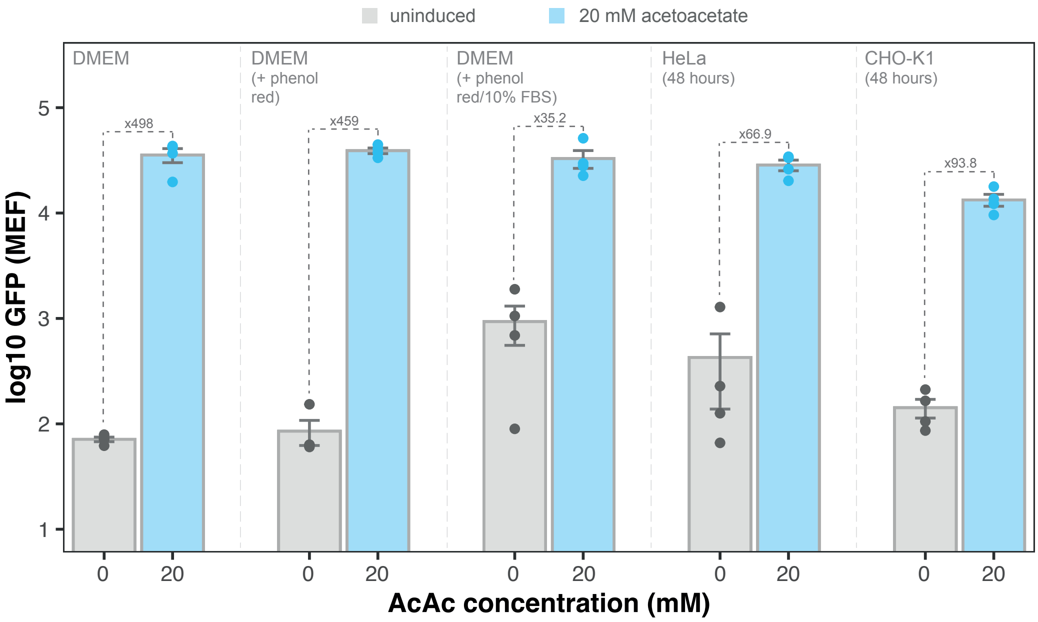


Figure S6: Normalised induction of the BW25113 ASAH0 whole-cell biosensor, in complex medias: DMEM recording media, DMEM media supplemented with phenol red, DMEM media supplemented with phenol red and 10% FBS, DMEM media conditioned with HeLa cells for 48 hours and DMEM media conditioned with CHO-K1 cells for 48 hours. The annotated numbers represent the fold change (calculated as mean normalised GFP at 20 mM induction divided by mean normalised GFP at 0 mM induction). ($n = 4$ biological repeats, points give medians and bars means of medians $\pm$ propagated SE)

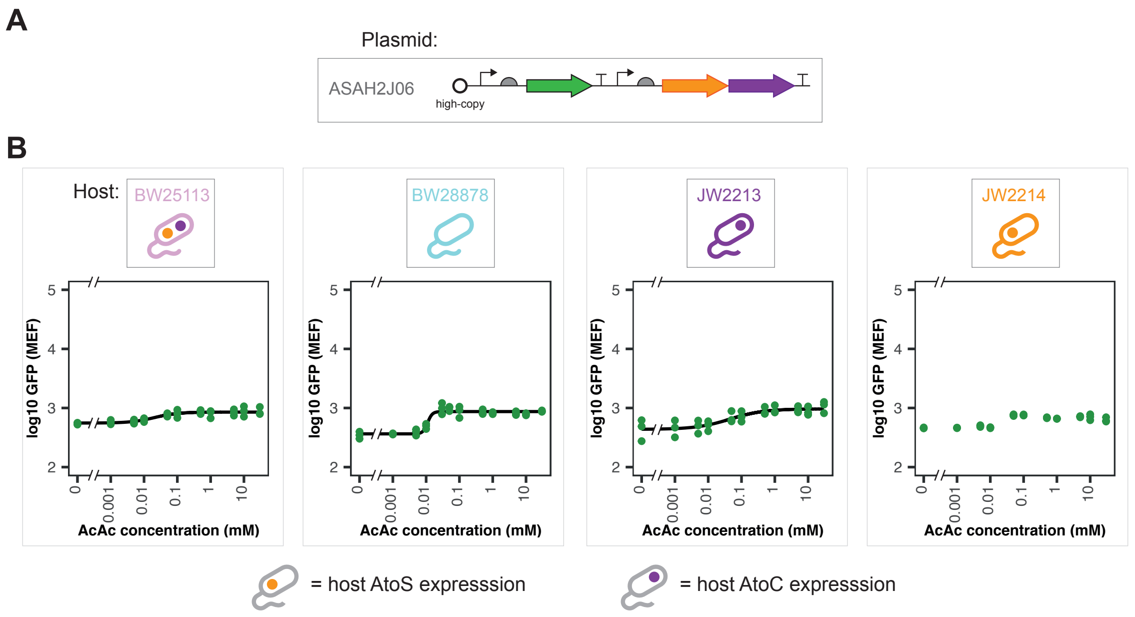


Figure S7: The performance of the ASAH2J06 biosensor plasmid in the four 'Ato' knockout strains. (**A**) Plasmid layout of ASAH2J06. (**B**) Median GFP fluorescence of ASAH2J06 plasmid in each of the given host chassis, at varying acetoacetate concentrations. (n = 3 biological repeats, data fit with Hill function)

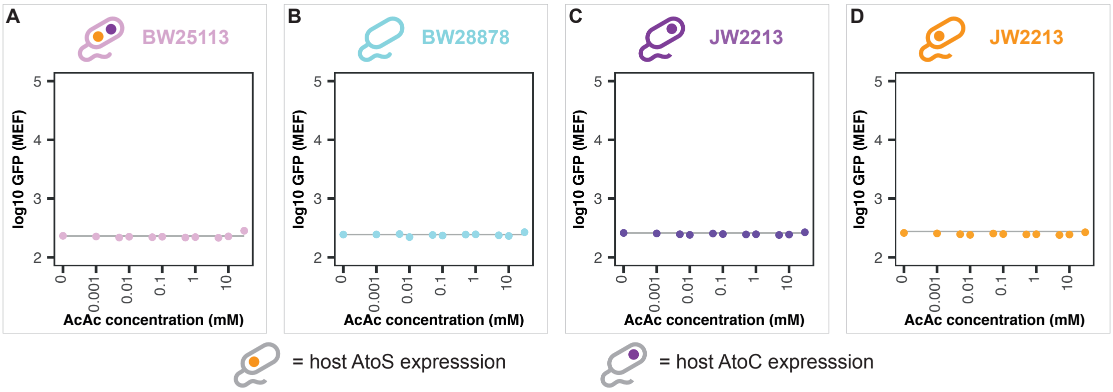


Figure S8: Concentration assays of the four empty 'Ato' knockout host strains. Strains: (**A**) BW25113, (**B**) BW28878, (**C**) JW2213 and (**D**) JW2214. Points give medians, lines show median GFP fluorescence of uninduced cells. (n = 1 biological repeat)

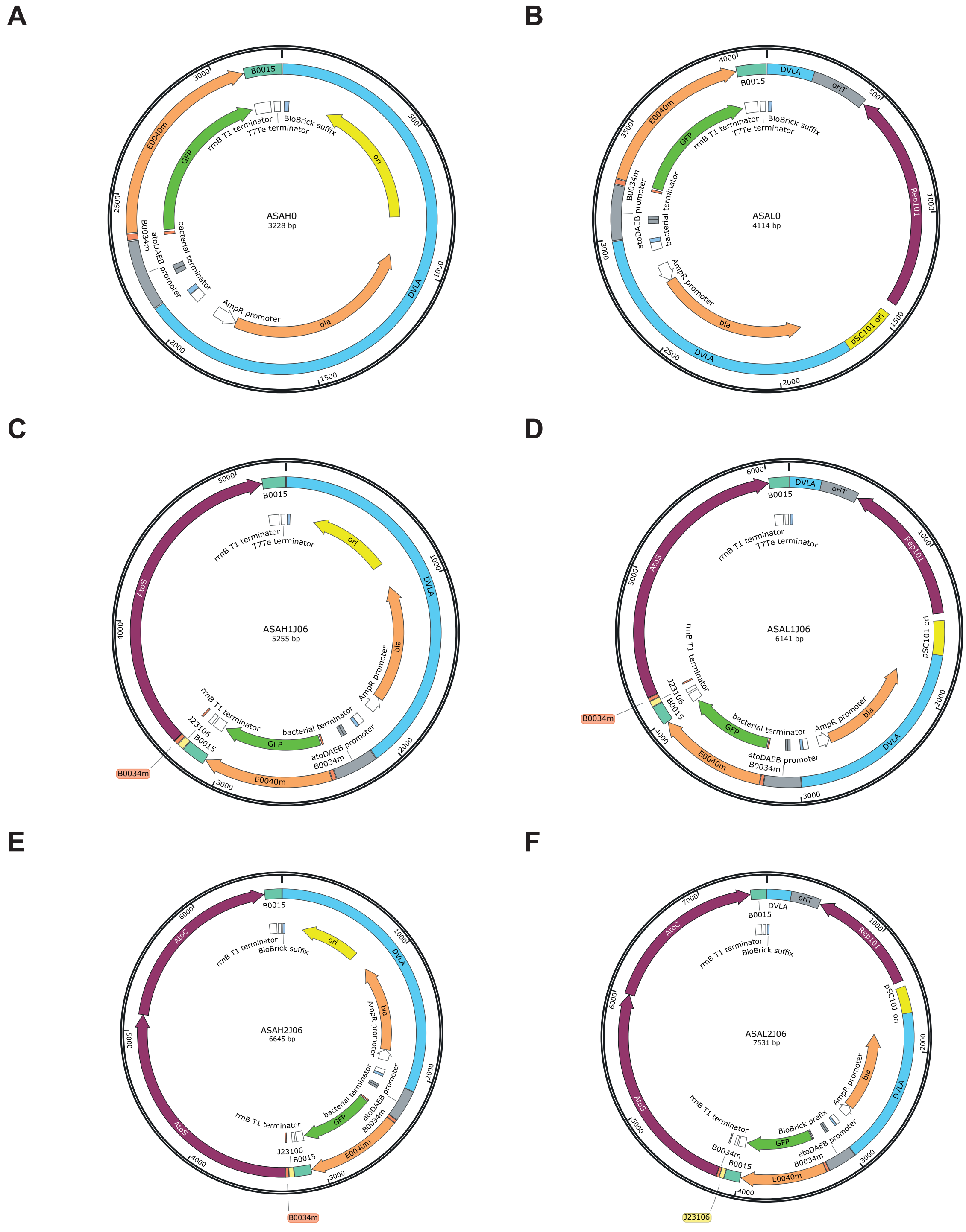


Figure S9: Plasmid maps of the six biosensor plasmids characterised within the main text: (**A**) high-copy ASAH0, (**B**) low-copy ASALO, (**C**) high-copy, *atoS*-expressing ASAH1J06, (**D**) low-copy, *atoS*-expressing ASAL1J06, (**E**) high-copy, *atoSC*-expressing ASAH2J06 and (**F**) low-copy, *atoSC*-expressing ASAL2J06.
